# Dynamic Modeling and Stochastic Simulation of Metabolic Networks

**DOI:** 10.1101/336677

**Authors:** Emalie J. Clement, Ghada A. Soliman, Beata J. Wysocki, Paul H. Davis, Tadeusz A. Wysocki

## Abstract

Increased technological methods have enabled the investigation of biology at nanoscale levels. Nevertheless, such systems necessitate the use of computational methods to comprehend the complex interactions occurring. Traditionally, dynamics of metabolic systems are described by ordinary differential equations producing a deterministic result which neglects the intrinsic heterogeneity of biological systems. More recently, stochastic modeling approaches have gained popularity with the capacity to provide more realistic outcomes. Yet, solving stochastic algorithms tend to be computationally intensive processes. Employing the queueing theory, an approach commonly used to evaluate telecommunication networks, reduces the computational power required to generate simulated results, while simultaneously reducing expansion of errors inherent to classical deterministic approaches. Herein, we present the application of queueing theory to efficiently simulate stochastic metabolic networks. For the current model, we utilize glycolysis to demonstrate the power of the proposed modeling methods, and we describe simulation and pharmacological inhibition in glycolysis to further exemplify modeling capabilities.

**Author Summary:** Computational biology is increasingly used to understand biological occurances and complex dynamics. Biological modeling, in general, aims to represent a biological system with computational approaches, as realistically and accurate as current methods allow. Metabolomics and metabolic systems have emerged as an important aspect of cellular biology, allowing a more sentive view for understanding the complex interactions occurring intracellularly as a result of normal or perturbed (or diseased) states. To understand metabolic changes, many researchers have commonly used Ordianary Differential Equations to produce *in silico* models of the *in vitro* system of interest. While these have been beneficial to date, continuing to advance computational methods of analyzing such systems is of interest. Stochastic models that include randomness have been known to produce more reaslistic results, yet the difficulty and intesive time component urges additional methods and techniques to be developed. In the present research, we propose using queueing networks as a technique to model complex metabolic systems, doing such with a model of glycolysis, a core metabolic pathway.

## Introduction

Cellular metabolism is an extensively complex network of enzymes, metabolites and other biomolecules required to both maintain homeostasis and appropriately react to stimuli. Biochemists began examining cell metabolism in the mid-19th century, and with our advancement in both experimental techniques and computational capacity, increasing comprehension of metabolic intricacies has been realized. As a relatively new field, metabolomics studies are concerned with the detection and quantification of metabolites. When considering the complexity of the metabolome, the analytical side of metabolomics and metabolism can easily become daunting. The KEGG Compound database currently contains 18,047 metabolites and other small molecules, making the intuitive aspect of understanding metabolite dynamics nearly inconceivable [1]. Thus, computational modeling reconstruction and simulation of metabolic systems have become pivotal in the analysis and surveillance of such systems.

Within the last decade, it has been a considerable goal to develop mathematical models of biological systems that accurately predict cellular and ultimately systems level behavior, providing quantitative details and prediction of phenotypic changes resulting from perturbation. Models as a whole are a representation of reality, aiming to accurately represent the system of study. Inclusion of all cellular components indirectly or directly involved are considered far too complex to model. Consequently, simplifications and assumptions must be made and often the perceived non-pivotal details, such as stochasticity, omitted. Nevertheless, the accuracy and competence of the model is dependent on these assumptions and simplifications.

Many approaches may be taken to model the dynamics of metabolic systems; importantly, the categorization of deterministic and stochastic modeling approaches. Most often, kinetic models of metabolism have been modeled using ordinary differential equations (ODE) providing a deterministic modeling approach that gives quantitative information on the interactions, underlying dynamics, and regulation of the components of the system [2]. ODE models operate with the assumption that all reactions occur under evenly mixed, homogenous populations with many molecules in the environment. From early on, ODEs have been used to simulate biochemical kinetics and biochemical networks. This approach, with historically “limited” computational power, was sufficient to describe the interactions and dynamics occurring within biochemical networks. Rapoport et al. describes the ability to determine metabolite concentrations of glycolytic intermediates in erythrocytes by a desktop calculator [3]. In our current time, with the aid of increasing computational power, metabolite concentrations within an enzymatic chain of reactions can be determined almost instantly. There are numerous methods and well defined strategies for solving ODEs; the prevalence and significance in both biochemical simulations in addition to mathematics offer a firm grasp on dealing with simple systems of ordinary differential equations. While ODE modeling reduces computational efforts, the assumptions and simplifications come at the cost of omitting noise and randomization that is inherent in biological systems. Thus, stochastic modeling approaches may be a more realistic representation of *in vitro* and *in vivo* systems [2].

Though ODE methods have been well defined in biological community, more recently, systems biology has begun to extend the limits of what has previously been capable computationally; modeling the complexities of biological variation - the stochastic effects inherent in biology. Stochastic models are typically formulated by the chemical master equation (CME), and have the ability to capture the stochastic occurrences common in biological systems. Yet, the drawback comes with the increased mathematical and computational complexity, additionally limiting the size of the network. The CME is a continuous Markov process that has commonly been handled by way of Monte-Carlo simulations, wherein the probability of a particular reaction occurring is calculated and the probability of the particular reaction occurring in a given time interval is also calculated and updated given the state of the system [4]. Needless to say, the complexity of even a small system becomes unmanageable. Not to mention the increase in the number of parameters required and whether or not the specific parameters are appropriate given the context in which they are derived [5]. The Gillespie algorithm, introduced in 1977, provides exact simulation methods and can be sped up with the implementation of Tau leaps [4,6,7]. Still, attempts are being made to improve stochastic simulation, and overcome the difficulties involved [2,5,8–13].

One relatively new approach to understanding and modeling complex biological networks is the application of queueing networks. Having some similarities to the Gillespie algorithm, queueing networks can be represented as a Markov chain, being more convenient to use than directly implementing the Gillespie algorithm [14]. Queueing networks have previously been used to describe data communication networks [15], servicing of patients at hospitals [16], the HIV infection process [17], pharmacokinetic modeling [18], and non-viral gene delivery [19]. Moreover, the implementation as described by Martin et al. [19], has accounted for other cell processes, like mitosis or cell necrosis, which are not easy to implement in the ODE approach. Queuing networks have also been used to develop a simple model of metabolism [20] and enzyme substrate interactions [21]. Briefly, queueing theory is a method of approaching stochastic simulations, doing so in such a way that computationally, it is less intensive and accordingly, possesses the ability to potentially describe larger networks - large networks that may not be able to be simulated in a reasonable amount of time given other stochastic simulation methods.

Recently, we have developed a tool to recapitulate observed *in vitro* insulin responses, plus measure the effects of Wortmannin-like inhibition on glucose uptake [22]. This has provided insight into transient changes in molecule concentrations throughout the insulin signaling pathway, and opened the door to identify critical, drug-targetable components of this pathway, including those associated with insulin resistance. Notably, our model was capable of calculating all network components of 100 averaged cells at near real-time: approximately 12 minutes on a desktop computer to produce 10 minutes of simulated data. Comparatively, the classical ODE approach of this complex network was calculated to take more than 80 days for completion. More importantly, though, the classical approach fails to take into account random variation naturally occurring within cells and tissues [23]. Conversely, the application of queueing theory has recently provided the ability to overcome computational intensity and incorporate a natural variation of kinetic constants and initial molecular concentrations. Herein, we present the current model using queueing theory to simulate the stochastic effects of glucose metabolism as a simplified technique to model metabolism, specifically glucose metabolism. For model validation, we provide qualitative comparisons of pharmacological inhibition in both simulated conditions and experimental metabolic data from cancer cells.

## Methods

The model presented uses ODE’s and glucose metabolism as a platform to describe the dynamical behavior of the pathway. Glucose metabolism has been described with ODEs in many modeling efforts. Being well defined both computationally and biochemically, researchers often model glycolysis and glucose metabolism when developing novel simulation methods. With a predetermined notion of the outcome, one can more easily compare results and begin validating the computational methods being developed. Consequently, we have used glycolysis to present queueing theory modeling of metabolic networks. A brief overview of glycolysis and glucose metabolism can be found in [24].

For the initial development of the model, we made use of previously derived mechanistic equations employing Michaelis-Menten kinetics. The mathematical analysis of the rate equations and parameters used are described in Mulukutla et al [25]. The derivation of the rate equations can be found in Mulquiney et al.. Our aim was to use a previously defined model to implement our proposed queueing theory approach, thus the parameters and kinetic constants for the current model were chosen to reflect the model investigated in Mulukutla et al. Notably, Mulquiney et al. report experimental and observed initial metabolite concentrations which were used for the current model. Concentration and parameter values that were either absent or substantially different between sources were obtained through additional literature searches. Furthermore, energy nucleotides and metal ions were fixed in our model for simplification and to centralize the model around the intermediate metabolites of glycolysis. Table 1 lists the initial concentrations of the metabolites measured in the simulation output, while Table 2 provides concentrations of additional substrates that were required for calculations, but not directly measured throughout the simulation.

**Table 1.** Initial Concentrations of Glycolytic Metabolites

| Metabolite | Concentration(mM) | Reference |
| --- | --- | --- |
| GLC | 5.0 | [1] |
| G6P | 0.039 | [1] |
| F6P | 0.013 | [1] |
| F1,6BP | 0.00231 | [1] |
| F2,6BP | 0.004 | [3] |
| DHAP | 0.02 | [2] |
| GAP | 0.00194 | [2] |
| 1,3BPG | 0.000369 | [1] |
| 3PG | 0.069 | [1] |
| 2PG | 0.01 | [1] |
| PEP | 0.017 | [1] |
| PYR | 0.0586 | [1] |

Intracellular concentrations for each metabolic intermediate. The metabolite concentrations (millimolar) are used in each simulation to initiate the model and are allowed to change over the course of the simulation.

**Table 2.**
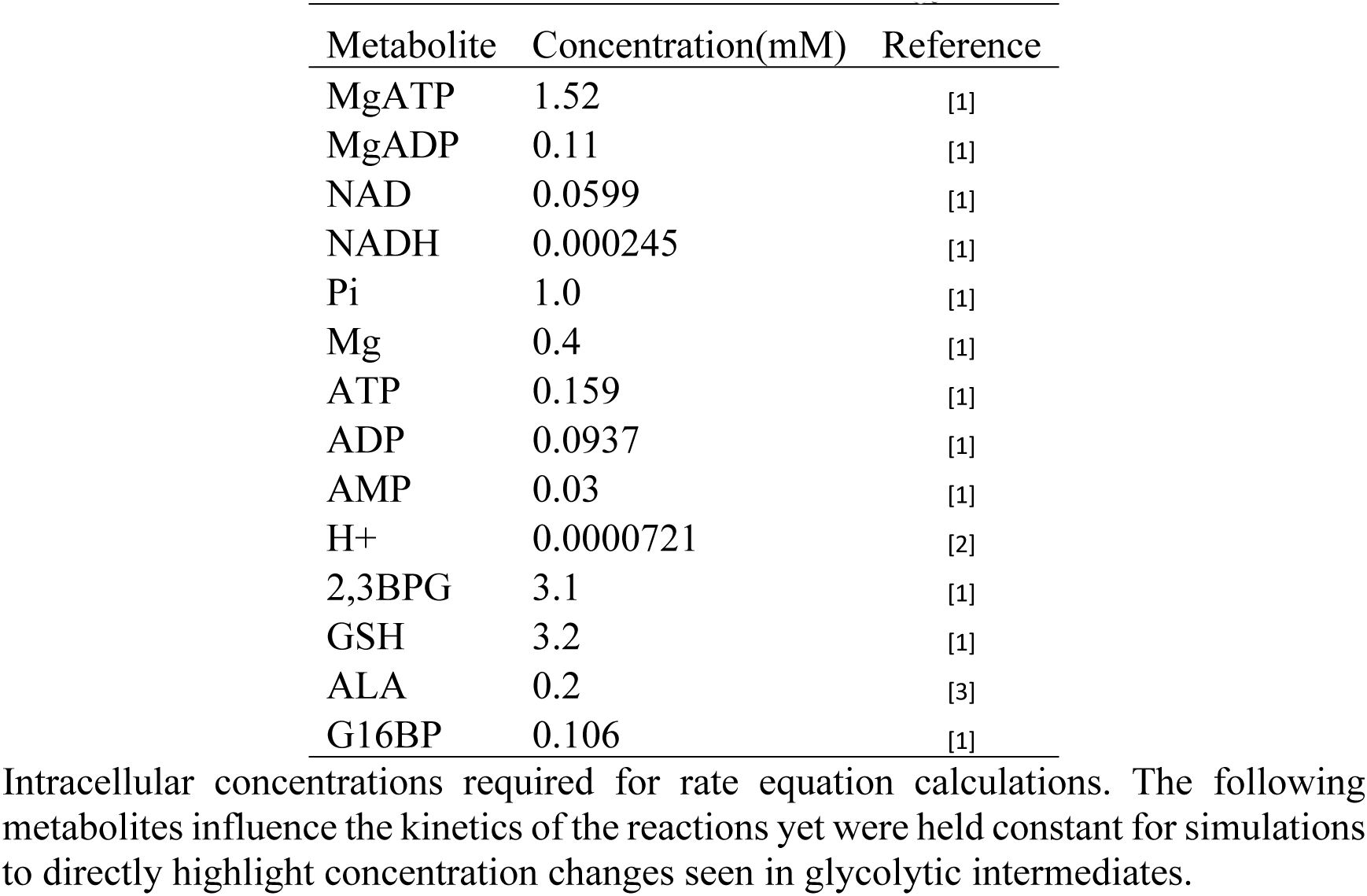
Additional Metabolites and Energy Nucle otides

| Metabolite | Concentration(mM) | Reference |
| --- | --- | --- |
| MgATP | 1.52 | [1] |
| MgADP | 0.11 | [1] |
| NAD | 0.0599 | [1] |
| NADH | 0.000245 | [1] |
| Pi | 1.0 | [1] |
| Mg | 0.4 | [1] |
| ATP | 0.159 | [1] |
| ADP | 0.0937 | [1] |
| AMP | 0.03 | [1] |
| H+ | 0.0000721 | [2] |
| 2,3BPG | 3.1 | [1] |
| GSH | 3.2 | [1] |
| ALA | 0.2 | [3] |
| G16BP | 0.106 | [1] |

Intracellular concentrations required for rate equation calculations. The following metabolites influence the kinetics of the reactions yet were held constant for simulations to directly highlight concentration changes seen in glycolytic intermediates.

To demonstrate the mechanics of how queueing networks are applied to modeling metabolic pathways, one can consider a pathway of *N* interacting metabolites *M*_1_, …, *M*_*N*_ having initial concentrations at time instant *t*_0_ of *C*_1_(*t*_0_), …, *C*_*N*_(*t*_0_). Within the considered metabolic pathway, each of the metabolites *M*_1_, …, *M*_*N*_ is involved in *K*_*i*_ reactions, *i* = 1, …, *N*. The corresponding reaction rates *v*_*i,j*_(*C*_1_(*t*), …, *C*_*N*_(*t*), *t*); *i* = 1, …, *N*; *j* = 1, …, *K*_*i*_, usually depend on the instantaneous concentrations of the interacting metabolites at time *t*, as well as other metabolites/enzymes, which given the time variability of those, is denoted as additional time dependability of *t*. The reaction rates can be positive or negative, with metabolites being produced for the positive sign and absorbed if the sign is negative. To find the concentration of the particular metabolite, *C*_*l*_(*t*) at given time instant *t*, one normally needs to solve a set of ODEs of the form:

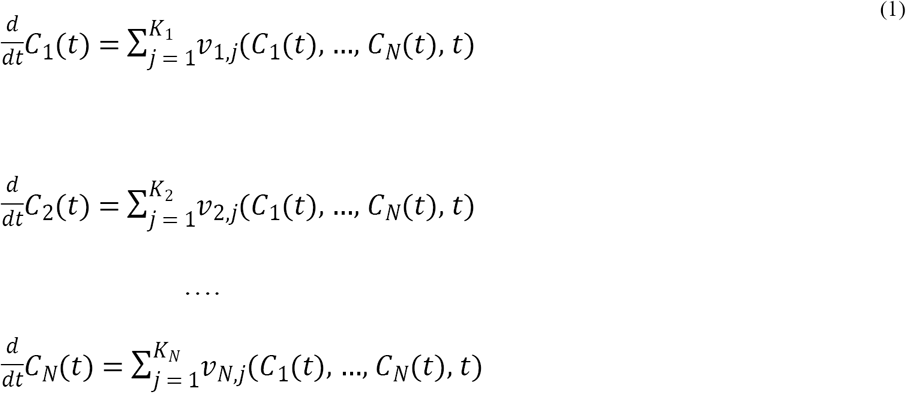

with the initial condition *C*_1_(*t*_0_), …, *C*_*N*_(*t*_0_).

Given the interdependency of the concentrations *C*_1_(*t*_0_), …, *C*_*N*_(*t*_0_), which is usually highly non-linear, and further dependency of other time varying and/or random factors, achieving the solution of such sets of equations is not only extremely computationally intensive, but also not guaranteed to produce a numerically stable result. The problem is further complicated by the fact that the concentrations *C*_1_(*t*), …, *C*_*N*_(*t*) are always non-negative, and as reported by Infante et al [30], and Erbe et al.[31], this is a non-trivial task or such a solution may not even exist. One can force the numerical solver to produce the non-negative solution, for example by using MATLAB’s ‘NonNegative’ option is in the ‘odeset’ solver [32]. However, this significantly increases the computation time, and the solution may not be accurate or numerically stable. This might be especially problematic for those metabolites that are not expressed in high concentrations and/or are very rapidly used in other reactions, for example, the metabolite glucose-6-phosphate (G6P) is also an intermediate in the Pentose Phosphate Pathway (PPP) and glycogen metabolism [33].

To find a method to simulate processes described by the set of ODEs (1), one can notice that each of the ODEs in (1) is of the form describing an average behavior of an *M*(*t*)/*M*(*t*)/*c* non-depleting queue [14]. In general, the *M*(*t*)/*M*(*t*)/*c* queue is such a system where arrivals form a single queue and are governed by a time varying Poisson process, there are *c* servers and job service times are exponentially distributed with time varying rates. The *M*(*t*)/*M*(*t*)/*c* non-depleting queues are special cases of queues [16] where for each time interval, the difference between corresponding arrival rate and service rate is non-negative. Massey et al [14] also analyzed a general case of *M*(*t*)/*M*(*t*)/*c* queues, for which there is no simple method to describe them by means of ODEs, but which can be depleted to zero elements in the queue or in other words for queues that can be completely emptied.

Hence, the *M*(*t*)/*M*(*t*)/*c* queues can be used to model metabolic pathways for simulation purposes, and instead of solving a set of ODEs (1), simulate a network of interconnected *M*(*t*)/*M*(*t*)/*c* queues, provided that the concentrations *C*_1_(*t*), …, *C*_*N*_(*t*) are digitized. The arrival rates are for the queues and the service rates are the reaction rates *v*_*i,j*_(*C*_1_(*t*), …, *C*_*N*_(*t*), *t*) normalized to the duration of a single simulation time step Δ*t*_*i*_ and the concentration increment Δ(*C*_*i*_(*t*)), which denotes a finite change of *C*_*i*_(*t*) in time increment of Δ*t*_*i*_. The normalization is done according to the formula:

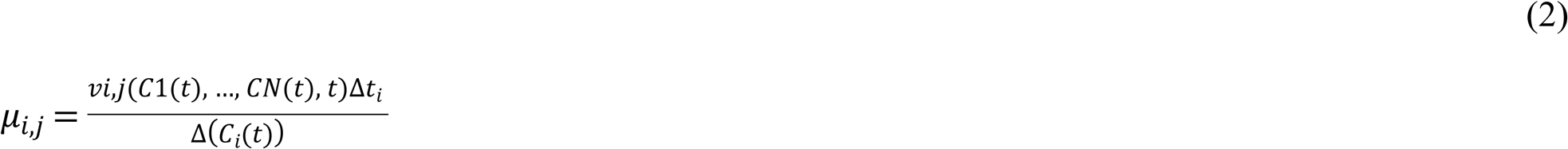

If the normalized rate *µ*_*i,j*_ is positive, it is an arrival rate while if it is negative it is a service rate. The instantaneous length of each queue provides a possible realization of a stochastic Markovian process representing variations of concentration of the given metabolite. Of course, the average changes in concentration can be achieved by averaging the simulation results for several simulation runs. To ensure correctness of simulation, the simulation time step Δ*t*_*i*_ and the concentration increment Δ(*C*_*i*_(*t*)), have to be chosen in such a way that all *µ*_*i,j*_ are lower than 1, as the arrival and service rates represent the probabilities of arrival and service of Δ(*C*_*i*_(*t*)) in the given time interval. It should be noted that for ensuring that just a single Δ(*C*_*i*_(*t*)) is processed in each time interval the conditions are as follows:

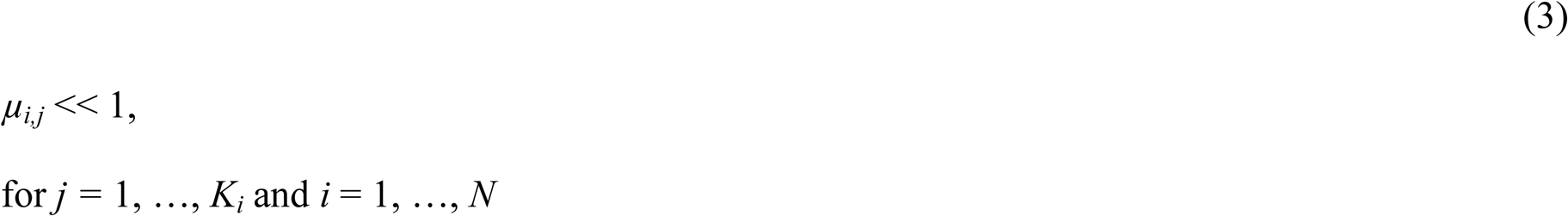

However, neither the simulation time step Δ*t*_*i*_ nor the concentration increment Δ(*C*_*i*_(*t*)), do not need to be the same for all *i* = 1, …,*N*, but can be chosen in a way that minimizes the simulation time, while ensuring that the condition (3) is satisfied. Though the time increment can be calculated dynamically within each step, for the current model we have chosen a constant time increment to speed up simulation time. Given the stochastic nature of chemical reactions, where the reaction rates can vary depending on environmental conditions, the reaction rates can be randomized by adding the Gaussian (or other) noise to the kinetic constants used to calculate values of *v*_*i,j*_(*C*_1_(*t*), …, *C*_*N*_(*t*), *t*). The same can be performed for the initial concentrations at time instant *t*_0_ of *C*_1_(*t*_0_), …, *C*_*N*_(*t*_0_).

A queue representing concentration of a single metabolite is shown in Fig 1, where the inputs to the represent reactions leading to production of the metabolite and outputs represent reactions using this metabolite. The cloud connected to the queue via bi-directional arrow represents processes not considered (or not even currently discovered) that result in an imbalance between an aggregated input to- and an aggregated output from the queue. The arrivals to the queue, representing discrete increments in concentration of the metabolite are modeled as Poisson processes, while the service time (time intervals between two consecutive output events) is modeled by an exponential distribution. Assuming that in total there are *c*-outputs from the queue, the queue can be considered as a standard *M*(*t*)/*M*(*t*)/*c* queue, as described before.

**Fig 1.**
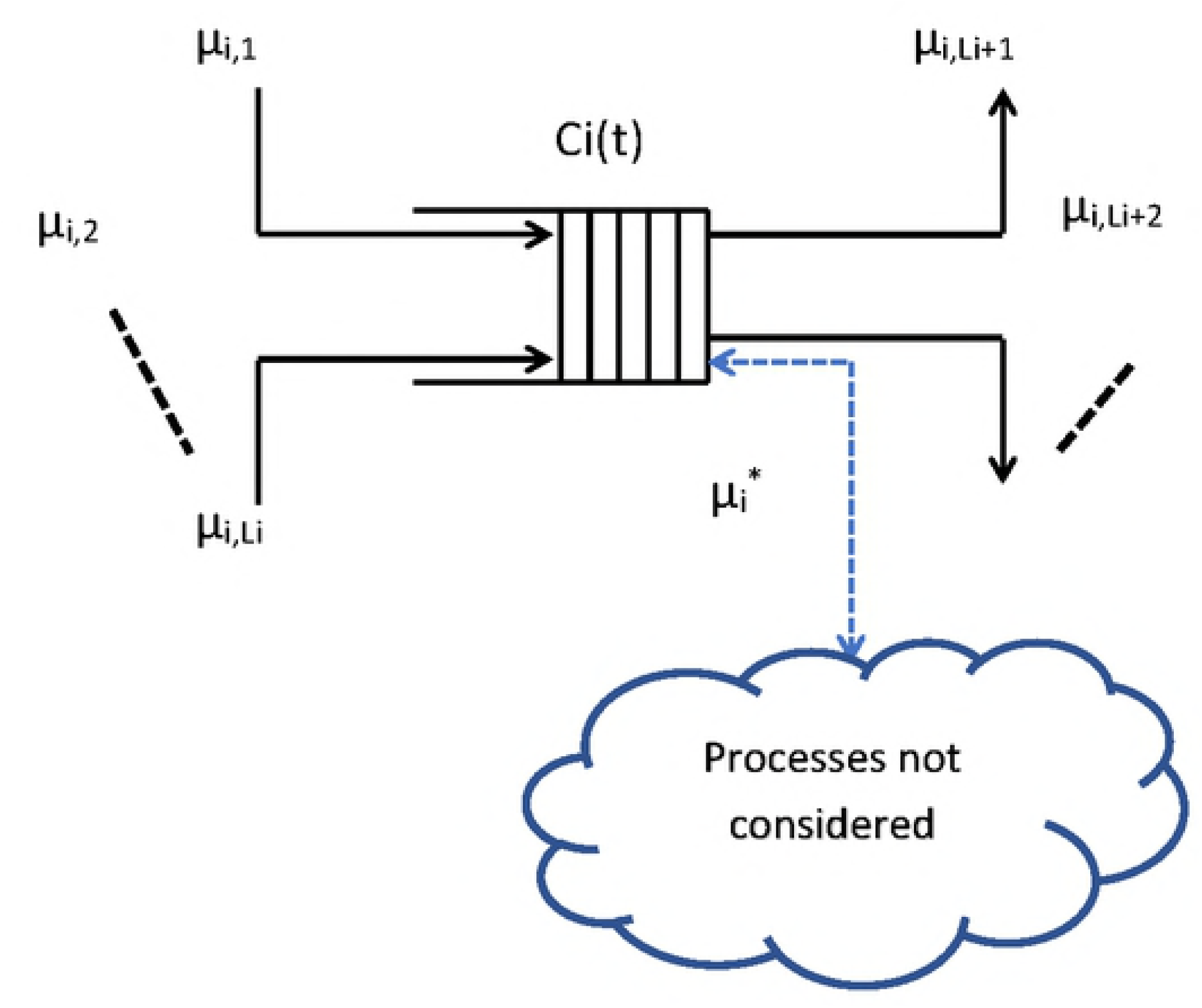
**Example Queue.** Queue representing concentration Ci(t) of the metabolite Mi; µi,j, j = 1, …, Li, are arrival rates as corresponding to processes resulting in production of metabolite Mi; µi,j, j = Li + 1, Li + 2, …, Ki*, are service rates corresponding to processes using metabolite Mi. Ki* = number of reactions considered in the model metabolite Mi is involved in.

For the description to be valid, the sums of all arrival rates, *µ*_*i,j*_, *j* = 1, …, *L*_*i*_, and the sum of all service rates 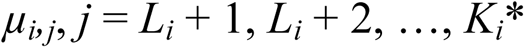 must each be lower than 1. A fulfilment of this condition can be satisfied by either reducing the duration of time increment or increasing the concentration unit. Of course, reducing the time increment increases the simulation time, as more simulation steps must be considered for the duration of an experiment, while increasing the concentration unit may reduce the accuracy of the simulation results. Therefore, a careful balance must be struck while choosing those parameters. Furthermore, from the perspective of implementing simulation of the metabolic process, it is convenient to ensure that in a given simulation step only one concentration unit of a given metabolite *M*_*i*_ is going to be processed. Assuming that there are 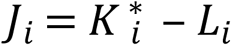 possible reactions that can be utilizing metabolite *M*_*i*_, the probability *P*_*i1*_ that this happens is given by the formula:

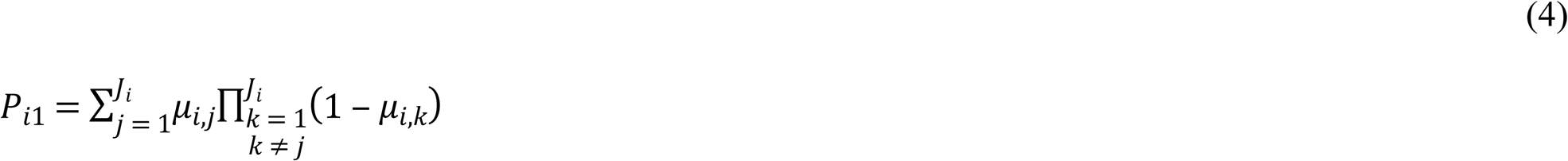

and the conditional probability *P*_*i*_{*j*|1} that if just one concentration unit is processed in a particular simulation step, it is processed in reaction associated with reaction rate *µ*_*i,j*_, is given by:

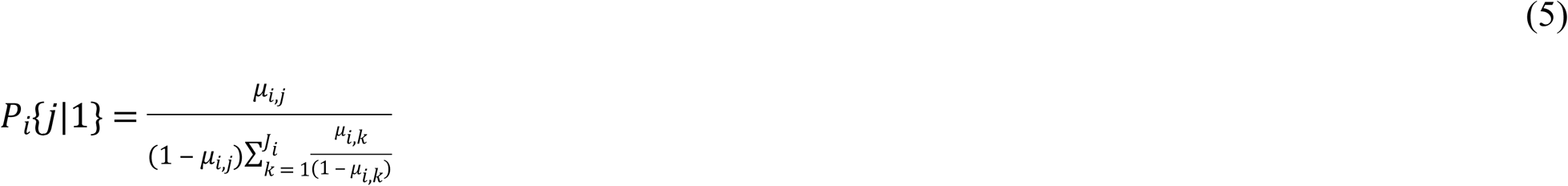

It is important to notice here that metabolomics data often include missing or semi-quantitative data, and some connections between the metabolites might have not been discovered, yet. To account for those unknown or missing reactions, an additional input/output pathway is included in the model for every metabolite considered, and shown in Fig 1 as a dashed line connection, which can be bi-directional. The rate *µ*_*i*_* is to be determined as the rate balancing the steady state value of the concentration *C*_*i*_(*t*). If the rate *µ*_*i*_* is positive, then for a corresponding metabolite concentration different from the steady state value, the rate is scaled by a factor equal to the ratio of the actual concentration to the concentration at the steady state. If it is negative, the scaling is inversely proportional. As previously mentioned, we have used queueing theory to describe additional biological pathways. In such, [34,35] provide further explanations of the proposed queueing theory methods. Additionally, pseudocode of the queueing theory application has been provided in the appendix section.

## Results

For the current study, our interest was in the ability to mechanistically model enzymatic reactions and stochastically simulate the dynamics of glycolysis utilizing queues. In general, queueing theory is a mathematical tool used to describe, model and analyze waiting lines, or queues [36]. At cellular level, metabolites are produced, absorbed, or used by cellular processes, thus forming “queues” of metabolites. Production or absorption of the metabolite adds to the appropriate queue length, and usage of the metabolite reduces the queue length. Both production and usage of a given metabolite are described by discrete random processes, referred to as an arrival and service process, respectively[37]. The queues can be easily interconnected, and as such are ideally suited to model metabolic networks, the same way as they are used to model the Internet [15]. Our previous work shows the ability to accurately simulate conditions seen *in vivo* using a fraction of the computing power of classical quantitative approaches at the time [22]. We have adjusted our queuing theory-based approached to model metabolic pathways given mechanistic rate equations of all glycolytic reactions and validated experimental metabolite data. As a core metabolic pathway common to all lifeforms, glycolysis is the enzymatic breakdown of glucose into a usable form of energy, additionally supplying intermediate metabolites as “building blocks” for connecting pathways to further support life. Naturally, glycolysis provides a scaffold to begin extending our model to incorporate additional metabolic pathways.

For the initial development of the model, we made use of previously derived mechanistic equations employing Michaelis-Menten kinetics. For the model simulations, all intermediate metabolites were represented by different queues, as described in the methods section. Additionally, the queues representing metabolites have been connected if there is a reaction converting one metabolite into another. Fig 2 shows the assembled queueing network representing glycolysis from glucose to pyruvate; where GLC, glucose; G6P, glucose 6-phosphate; F6P, fructose 6-phosphate; F16BP, fructose 1,6-bisphosphate; F26BP, fructose 2,6-bisphosphate; GAP glyceraldehyde 3-phosphate; DHAP, dihydroxyacetone phosphate; 13BPG, 1,3-bisphosphoglycerate; 3PG, 3-phosphoglycerate; 2PG, 2-phosphoglycerate; PEP, phosphoenolpyruvate; PYR, pyruvate.

**Fig 2.**
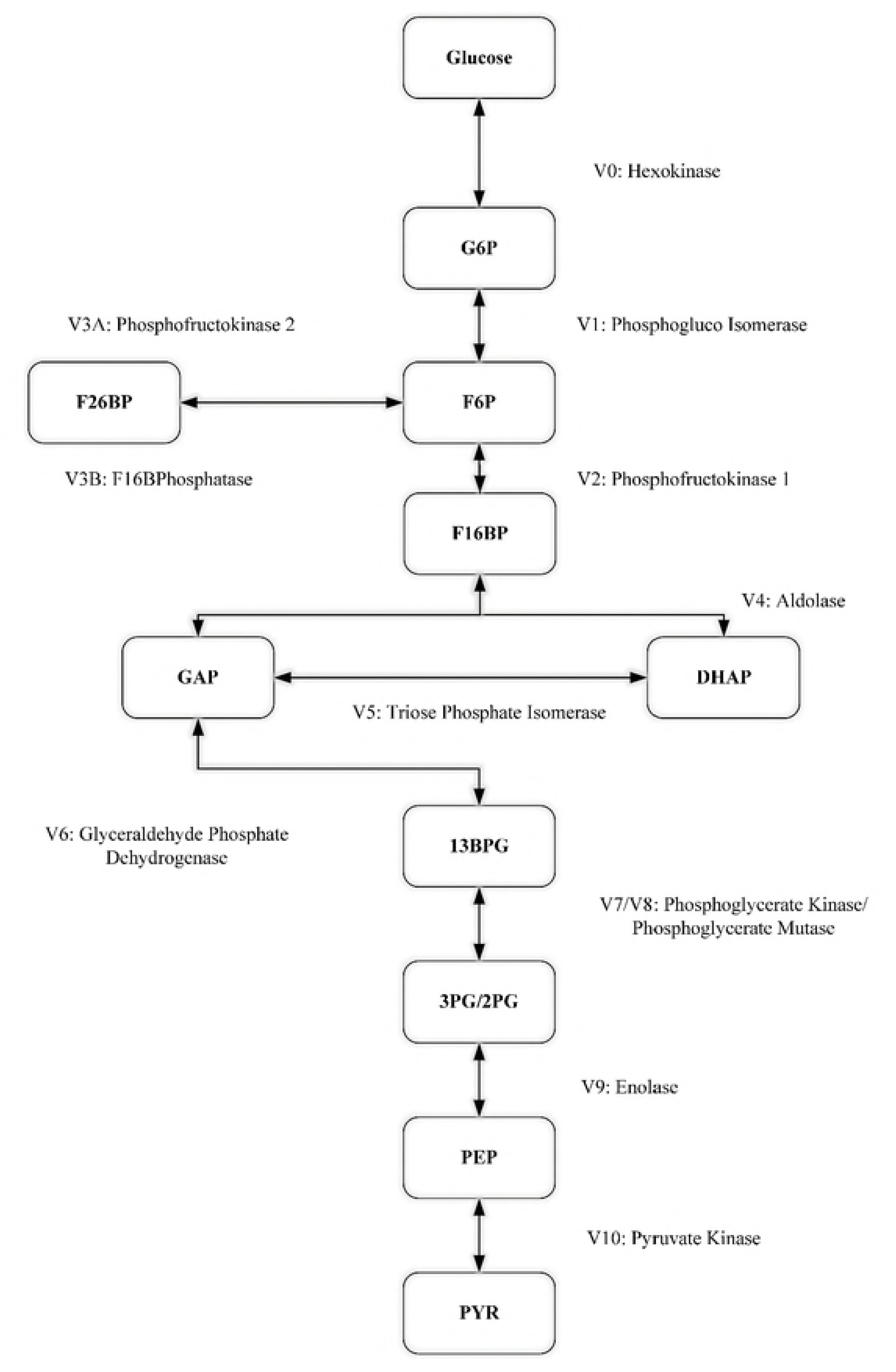
**Simulated metabolic pathway from glucose to pyruvate.** Arrows denote the modeled reactions. Vi, i = 0, …, 10, and V3A, V3B, are the reaction rates; for bidirectional arrows the direction is determined by the sign of the corresponding reaction rate with the positive direction being from the top down. GLC, glucose; G6P, glucose 6-phosphate; F6P, fructose 6-phosphate; F16BP, fructose 1,6-bisphosphate; F26BP, fructose 2,6-bisphosphate; GAP glyceraldehyde 3-phosphate; DHAP, dihydroxyacetone phosphate; 13BPG, 1,3-bisphosphoglycerate; 3PG, 3-phosphoglycerate; 2PG, 2-phosphoglycerate; PEP, phosphoenolpyruvate; PYR, pyruvate.

For the stochastic simulations presented, the rate equations and model parameters were used as they are indicated in the literature (Table 1 and Table 2). Highlighting the significance of the approach, the current methods enabled rapid alteration of parameter adjustment and additional simulations under a variety of selected *in silico* conditions. Due to the rapid catalytic conversion 3-PG and 2-PG in combination with the low metabolic concentrations, as separate queues the metabolites 3-PG and 2-PG were readily depleted given the 1 microsecond time scale used for the metabolite calculations. Thus, 3-PG and 2-PG were grouped in a single queue avoiding the occurrence of complete deficiency. Furthermore, the summation of the two consecutive metabolites within a queue slightly decreases total calculations and consequently, simulation time.

As previously noted biochemical reactions, and biological system, in general, are inherently stochastic processes. Consequently, randomness and variation were incorporated to add additional stochastic elements to the model simulations. Reaction rates were randomized during simulation by adding an arbitrarily chosen 10% Gaussian noise to the kinetic constants used to calculate values of vi,j(·). The same was performed for the initial concentrations at time instant t_0_ of all glycolytic intermediates. During the simulations, each simulated cell is calculated independently; that is, concentrations of each molecule in the metabolic network is stochastic, and bound by error values listed in the literature. The queueing theory approach causes the actual concentrations of given molecule types to be simulated as separate queues within each cell. The probability of a movement happening at any time slice from one queue to the next is determined by the relevant reaction speed. Movements between storages happen at a particular time instant if a number randomly drawn from the interval [0,1] at that time instant is smaller than the reaction speed governing the movement. After simulations have been performed for every considered cell, the results are averaged over the cell population. Variations of 10% glucose levels are randomly computed for every simulated second. The simulations were run with a 1 microsecond time step, and random variations in the values of kinetic constants used in calculating reaction rates were introduced every second. Initial concentrations were randomized by adding 10% Gaussian noise.

### Glycolytic Flux

Previously, Mulukutla et al. aimed to assess the control of different isoforms of the three rate limiting glycolytic enzymes on overall pathway flux behavior. The rate limiting enzymes of glycolysis, hexokinase (HK), phosphofructokinase (PFK), the bifunctional enzyme phophofructokinase-2/ fructose 2,6-bisphosphatase (PFKFB), and pyruvate kinase (PK) each have multiple isoforms and may be expressed in combination within single cell in a cell-type dependent manner. We considered regulatory mechanisms of PFK, PFKFB, and PK by including parameters and terms in the rate equations to consider the feedback inhibition and activation, keeping both upper and lower glycolytic regulatory loops active in our simulations. The feedback considered consists of F26BP (an important activator of glycolytic flux) and F16BP activation of PFK, F16BP activation of PK, and PEP inhibition of PFKFB activity. The parameters set to simulate the feedback loops are as follows: K_PFKf16bp=0.65 mM and K_PKf16bp=0.04 mM. The PFKFB kinase/phosphatase (K/P) ratio, the ratio between the kinase and phosphatase activity, was set to 0.1 by adjusting the value of the PFKFBPase Vmax leaving the kinase Vmax at its original value. Different K/P ratios are given in the literature based on specific tissue and cell type. The range varies from less than 1 to 710 depending on the isoform of PFKFB expressed and the tissue type in which it is found. Notably, PFKFB is highly dependent on signaling and hormonal regulation, which can transiently change the K/P ratio given the stimulus. Signaling regulation was not considered in this model, though this component is of interest for further study. Thus, we aimed to keep F26BP relatively constant throughout the initial steady state testing to keep the flux toward a stable level. We found that the K/P ratio of 0.1 kept F26BP and all other metabolites constant over time, the given the parameters used. Therefore, the 0.1 K/P ratio was used to further test the ability of the model to simulate metabolite changes. Simulations were repeated for 30 cells, once completed, the average concentrations of each metabolite per cell were graphed as a function of time (Fig 3).

**Fig 3.**
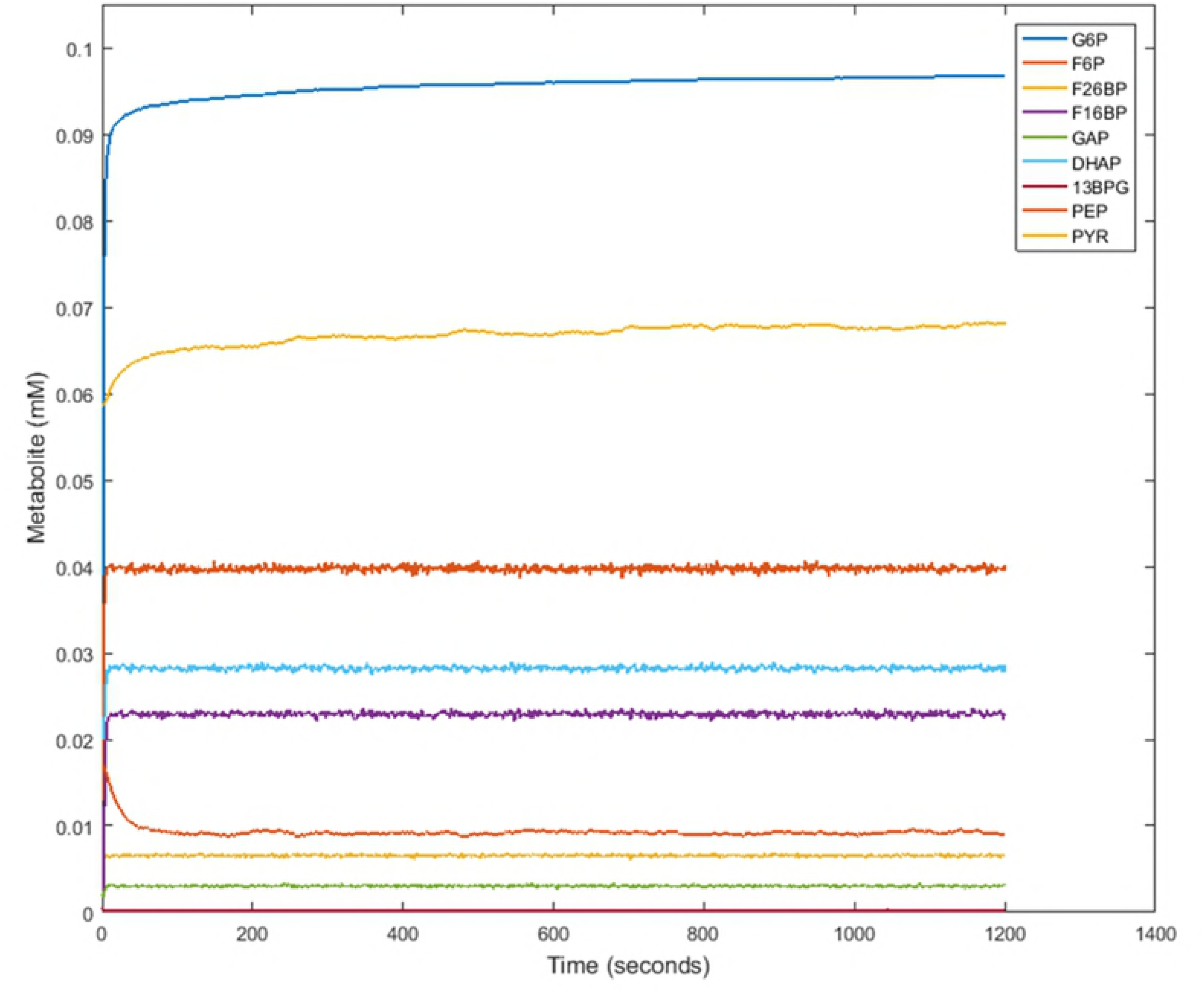
**Steady state glycolytic flux.** Metabolite concentrations were simulated with an input of 5 mM glucose over the course of 1200 seconds to model an unperturbed and constant state.

### GAPDH Inhibition

*In vitro* experiments and model simulations were performed to assess the performance of the proposed queueing approach. FK866 is a non-competitive inhibitor of Nicotinamide phosphoribosyl transferase (NAMPT), the enzyme that supplies the majority of the intracellular pool of NAD+, a required substrate for the GAPDH reaction. Extensive research has been performed analyzing the effects of FK866 on high glycolytic flux in cancer cells [38–41]. Under limited NAD+ concentrations, the GAPDH reaction represents a bottleneck in glycolysis producing a block in the glycolytic flux. Experimental results show the upper level glycolytic metabolites, including G6P, F6P, F16BP, GAP and DHAP accumulate while the lower metabolites, 13BPG, 3PG/2PG, PEP, and PYR, decrease as substrates become unavailable. Thus, we hypothesized that with the reduction of GAPDH activity and consequently simulation of enzyme inhibition *in silico*, the model should be able to mimic the qualitative metabolic trends seen *in vitro*. Notably, kinetics and enzyme concentrations for the specific cancer cell lines were unknown, therefore, to account for the differences between the cancerous and non-cancerous simulations, the reaction rates were scaled (See Supplemenary file S5). The effects of FK866 are presented in the experimental data provided by [40] in Figs 4, 6, 8 and 10, and by the present model outcomes of GAPDH activity inhibition in Figs 5, 7, 9 and 11.

**Fig 4.**
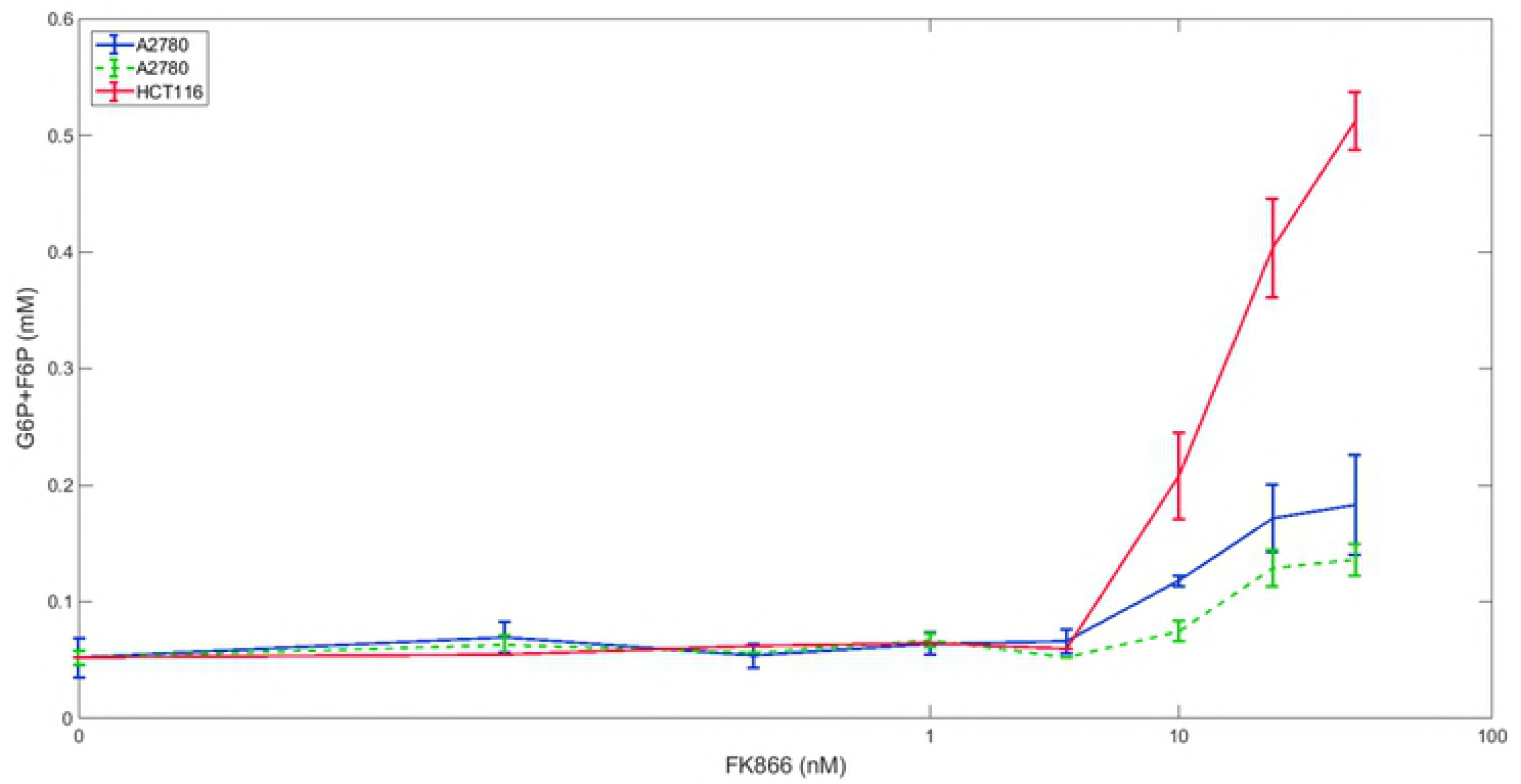
**Effects of FK866 on G6P and F6P concentrations in vitro.** Experimental metabolomics data measuring G6P and F6P concentrations with the inhibitor FK866 in (solid blue and dashed green lines) A2780 and (red) HCT116 cancer cells.

**Fig 5.**
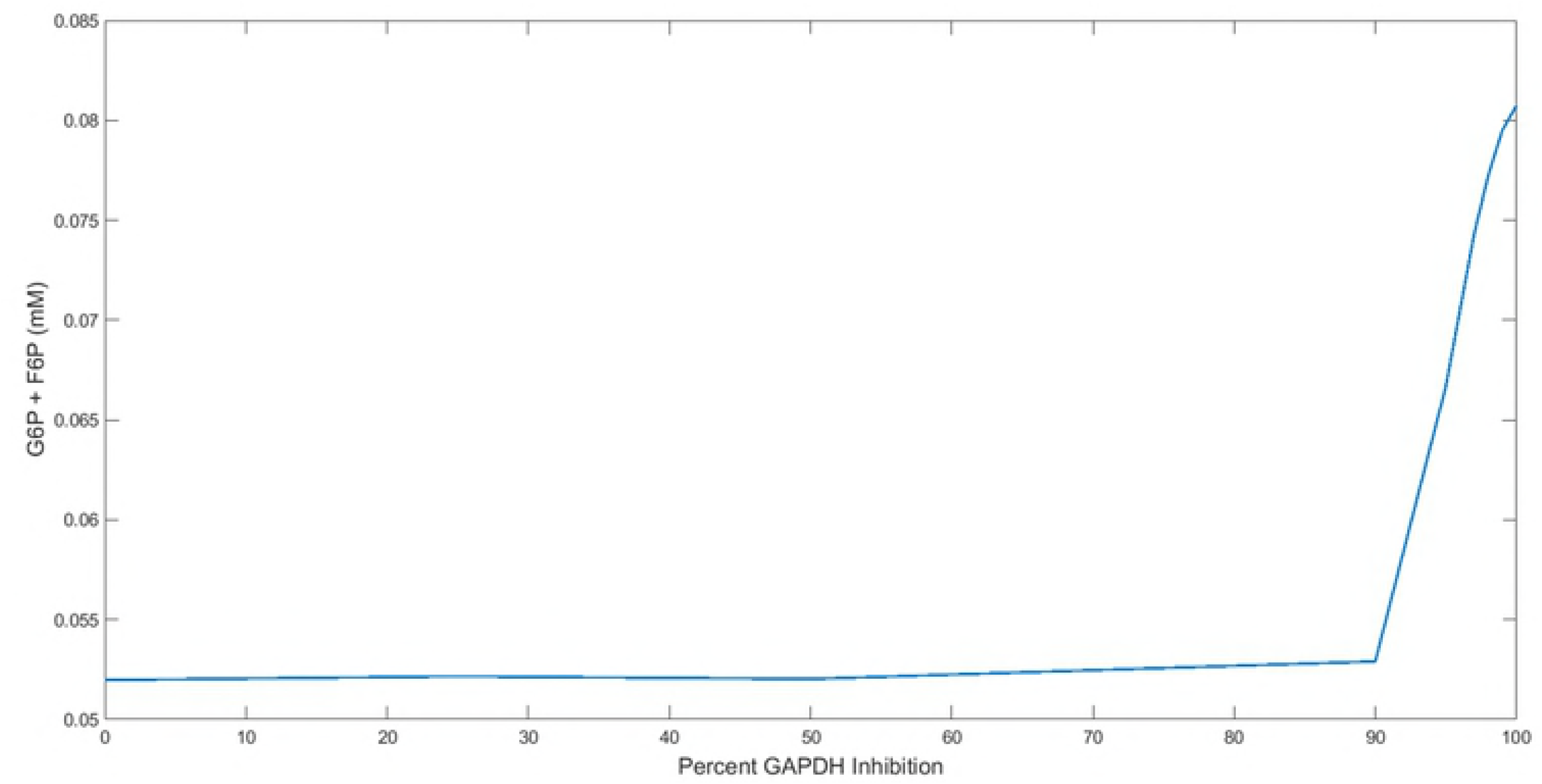
**Effects of GAPDH Inhibition on G6P and F6P concentrations in silico.** Simulation of the dynamics of glucose 6-phosphate (G6P) and fructose 6-phosphate (F6P) with the inhibition of Glyceraldehyde phosphate dehydrogenase (GAPDH). The Vmax of GAPDH was set at 0, 25, 50, 90, 95, 97, 98, 99, and 100 percent of its initial value in separate model simulations to simulate varied levels of pharmacological inhibition on the enzyme.

**Fig 6.**
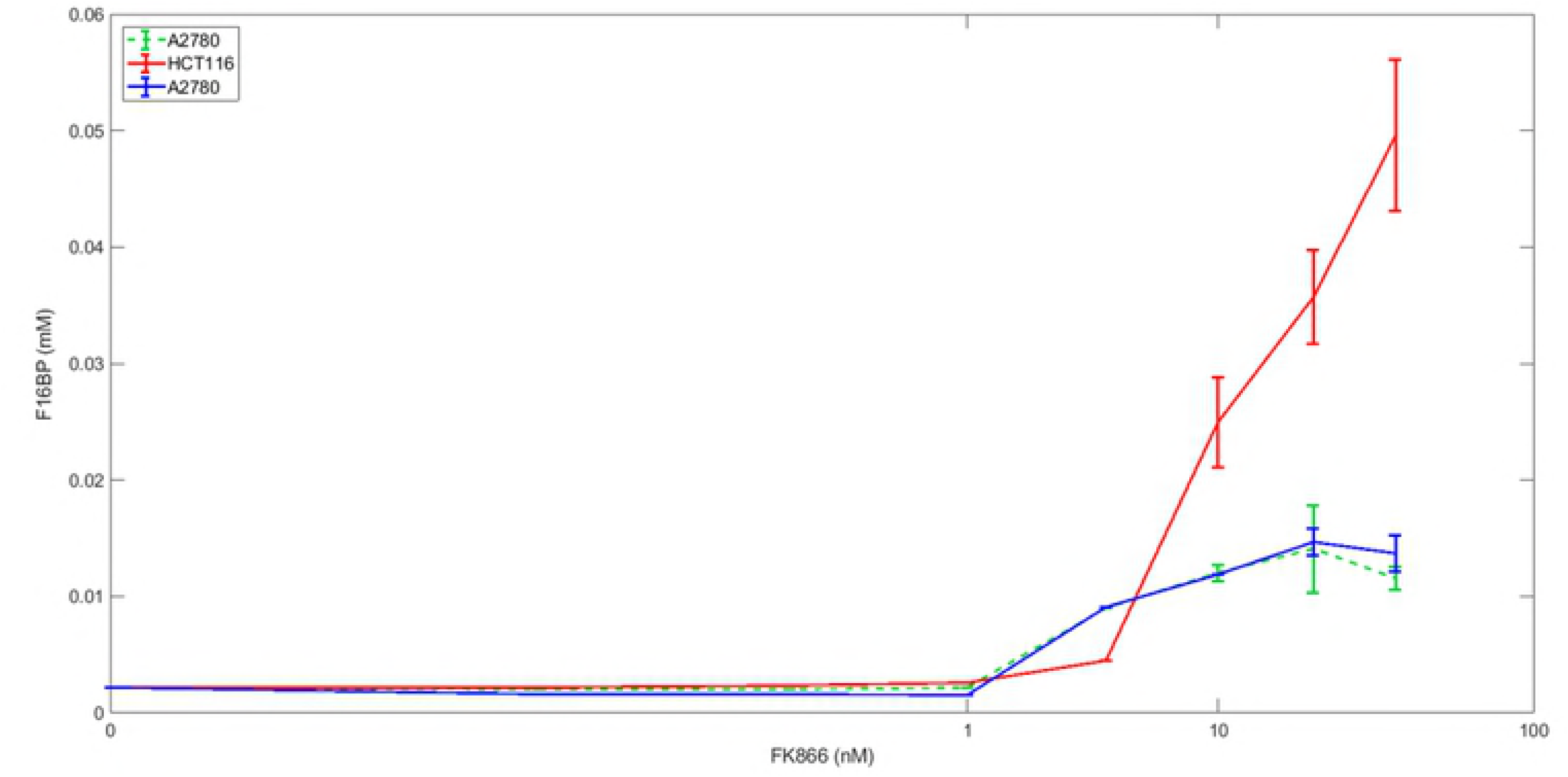
**Effects of FK866 on FBP concentrations in vitro.** Experimental metabolomics data measuring fructose 1,6-bisphosphate concentrations with the inhibitor FK866 in (solid blue and dashed green lines) A2780 and (red) HCT116 cancer cells.

**Fig 7.**
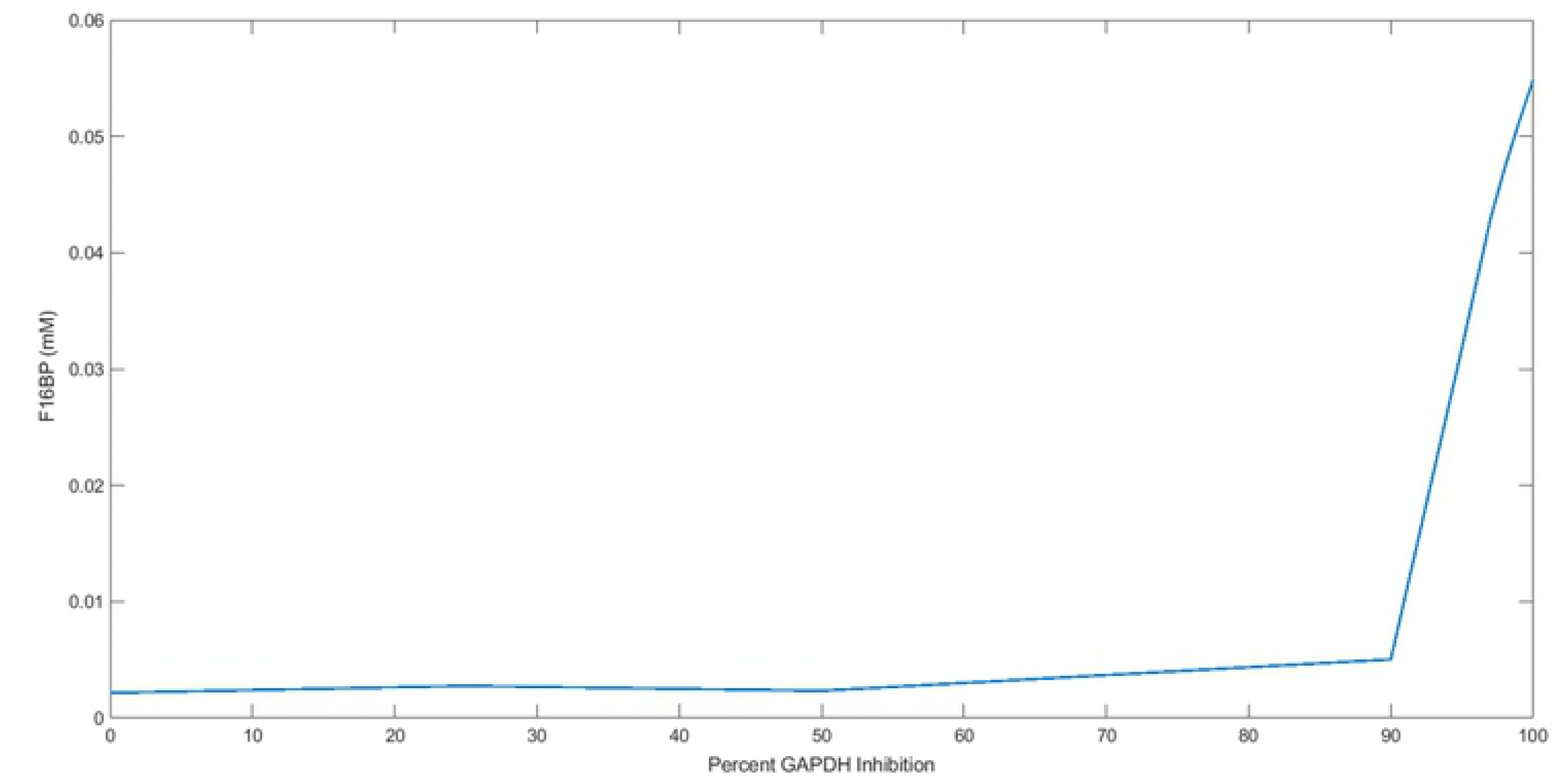
**Effects of GAPDH Inhibition on FBP concentrations in silico.** Simulation of the dynamics of fructose 1, 6-bisphosphate (FBP) with the inhibition of Glyceraldehyde phosphate dehydrogenase (GAPDH). The Vmax of GAPDH was set at 0, 25, 50, 90, 95, 97, 98, 99, and 100 percent of its initial value in separate model simulations to simulate varied levels of pharmacological inhibition on the enzyme.

**Fig 8.**
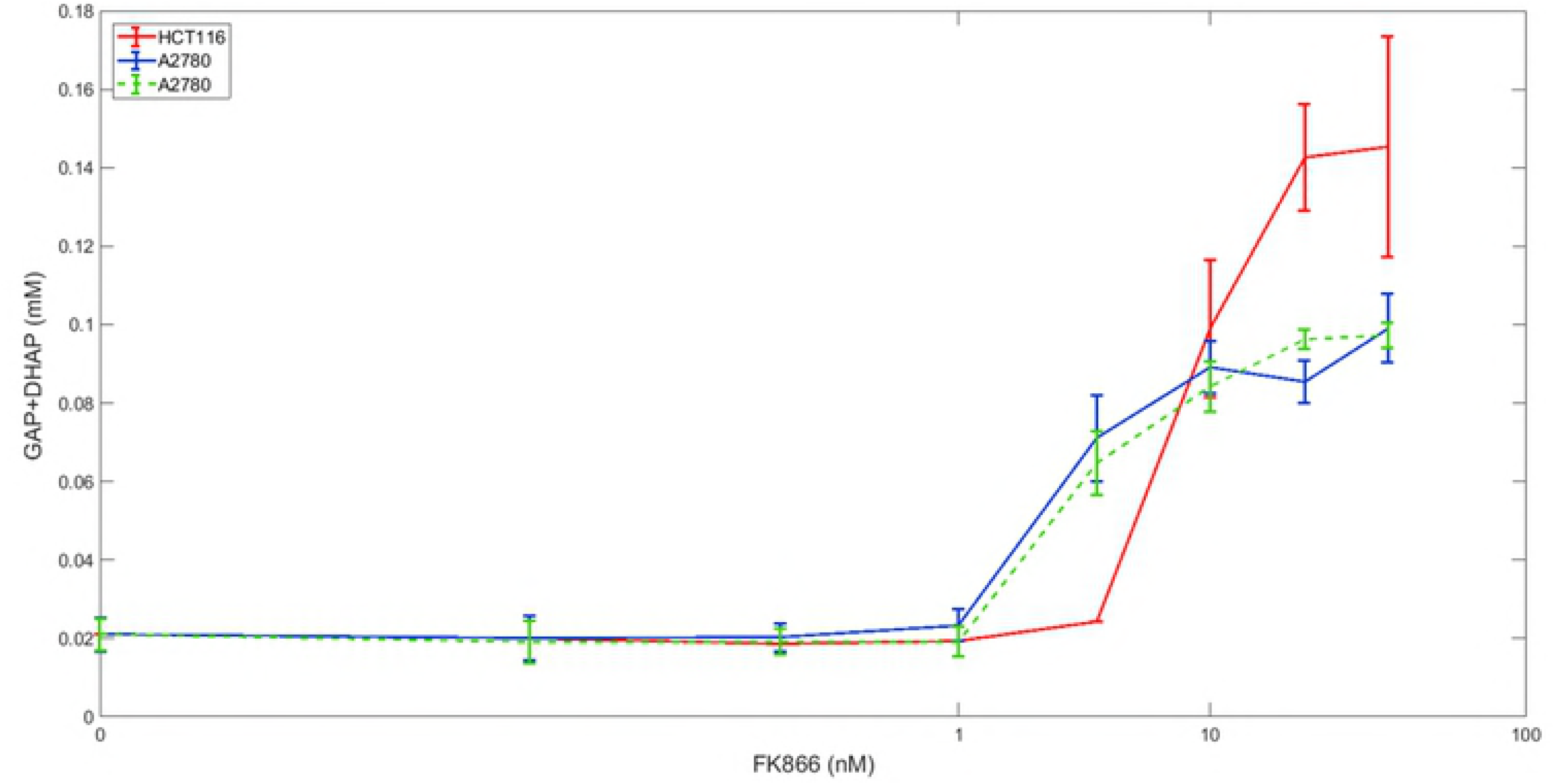
**Effects of FK866 on G6P and F6P concentrations in vitro.** Experimental metabolomics data measuring GAP and DHAP concentrations with the inhibitor FK866 in (solid blue and dashed green lines) A2780 and (red) HCT116 cancer cells.

**Fig 9.**
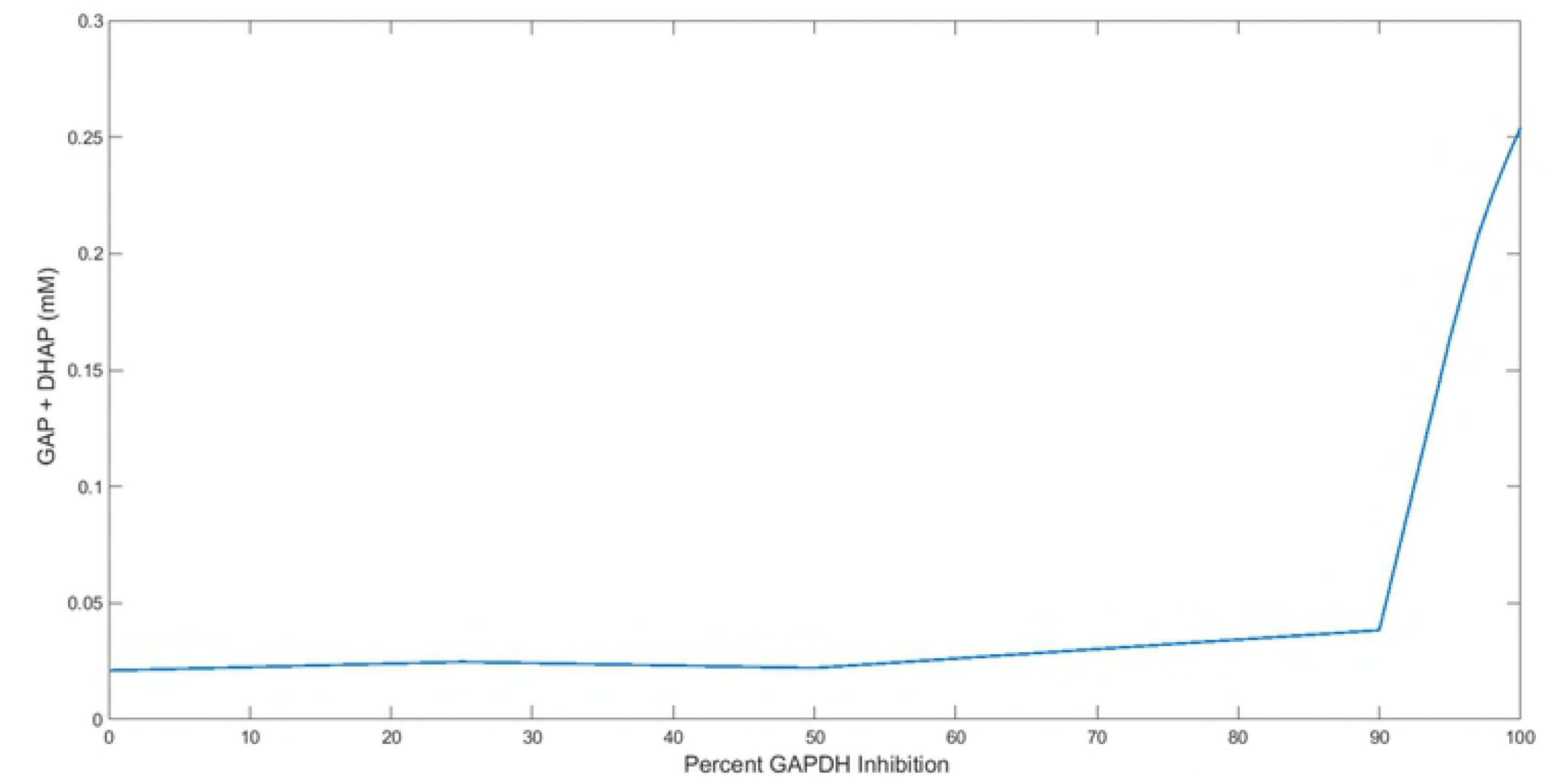
**Effects of GAPDH Inhibition on GAP and DHAP concentrations in silico.** Simulation of the dynamics of glyceraldehyde 3-phosphate (GAP) and dihydroxyacetone phosphate (DHAP) with the inhibition of Glyceraldehyde phosphate dehydrogenase (GAPDH). The Vmax of GAPDH was set at 0, 25, 50, 90, 95, 97, 98, 99, and 100 percent of its initial value in separate model simulations to simulate varied levels of pharmacological inhibition on the enzyme.

**Fig 10.**
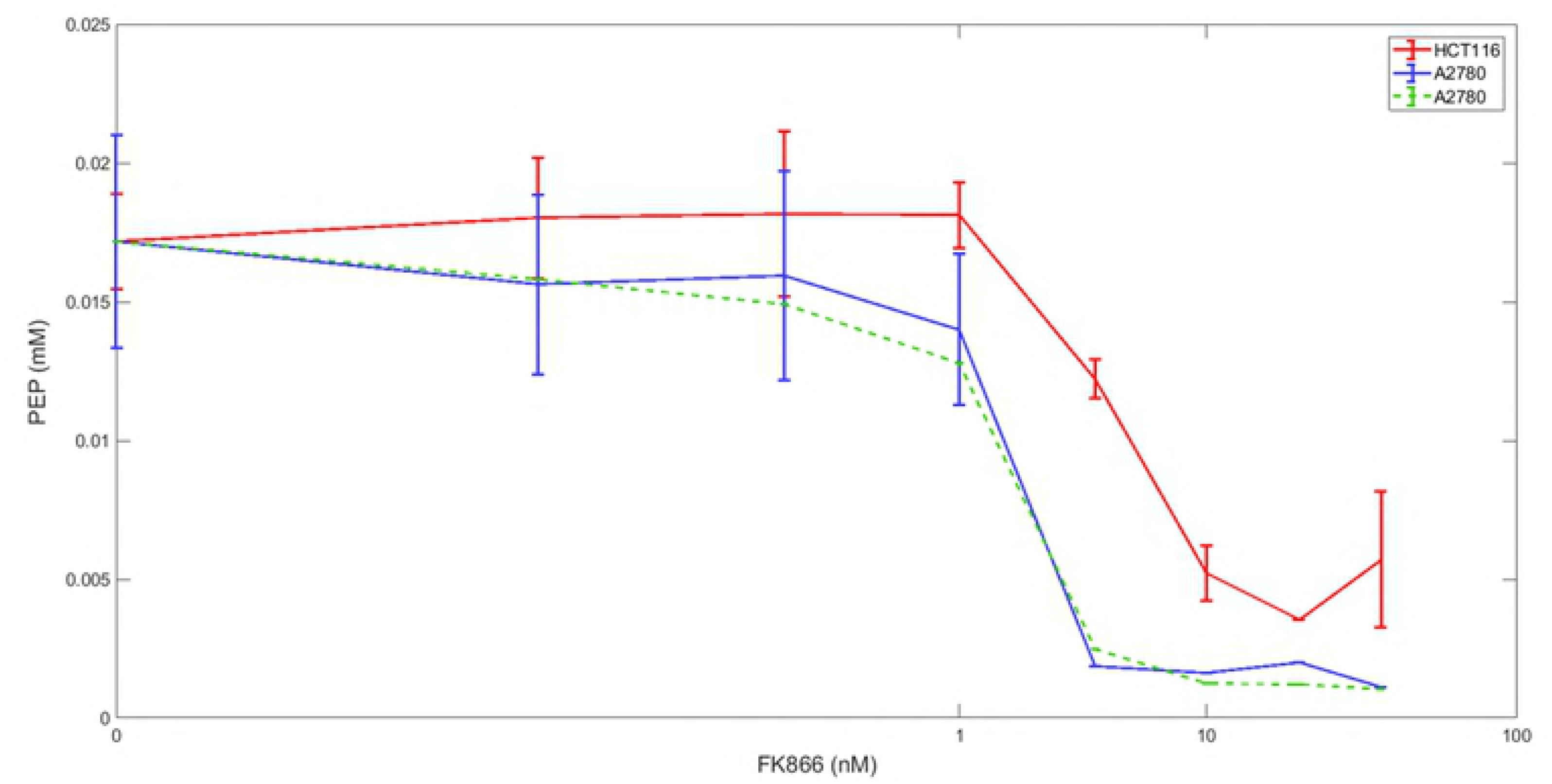
**Effects of FK866 on PEP concentrations in vitro.** Experimental metabolomics data measuring G6P and F6P concentrations with the inhibitor FK866 in (solid blue and dashed green lines) A2780 and (red) HCT116 cancer cells.

**Fig 11.**
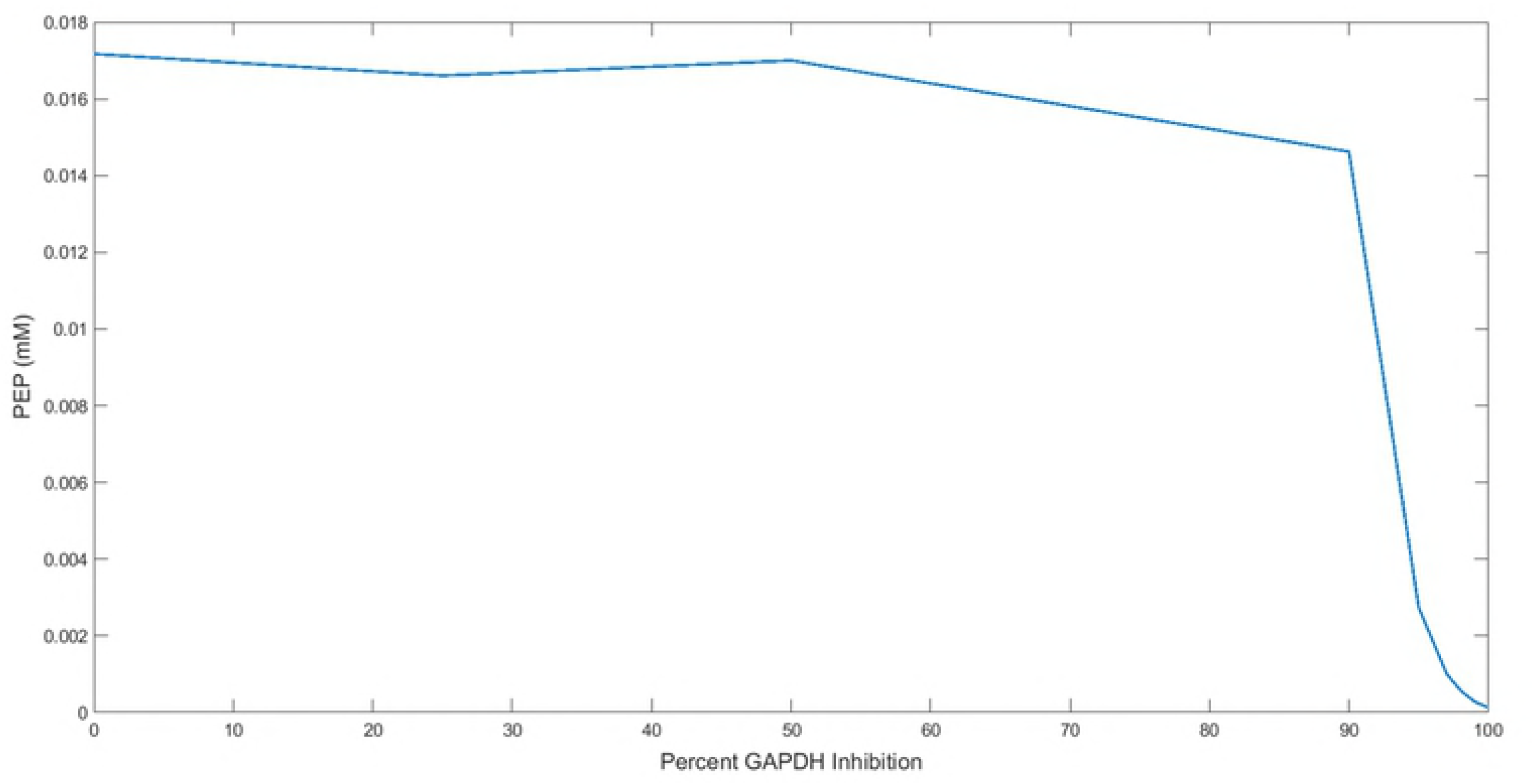
**Effects of GAPDH Inhibition on PEP concentrations in silico.** Simulation of the dynamics of phosphoenolpyruvate (PEP) with the inhibition of Glyceraldehyde phosphate dehydrogenase (GAPDH). The Vmax of GAPDH was set at 0, 25, 50, 90, 95, 97, 98, 99, and 100 percent of its initial value in separate model simulations to simulate varied levels of pharmacological inhibition on the enzyme.

GAPDH activity was reduced by adjusting the Vmax of the reaction catalyzed by GAPDH. Inhibition was simulated at 0%, 25%, 50%, 90%, 95%, 97%, 98%, 99%, 100% GAPDH activity, with each inhibitory level run as a separate simulation for a cell population of 30. The simulated results of GAPDH inhibition of G6P+F6P, F16BP, GAP+DHAP, and PEP, are plotted as dose reponse curves in Figs 5, 7, 9 and 11 to reproduce the effect of metabolite changes from experimental (Figs 4, 6, 8 and 10) pharmacological inhibition of two cancerous cell lines. The FK866 inhibitor concentrations used in both A2780 ovarian and HCT116 colorectal cancer cell lines *in vitro*, were compared to *in silico* reduction of the percent GAPDH activity. Of note, we are making the assumption that at the lowest FK866 concentration (0.3 nM) used *in vitro* did not inhibit GAPDH activity, whereas the highest inhibitor concentrations (40 nM) used completely inhibit its forward catalytic activity since the degree of inhibition corresponding to an exact inhibitor dosage is not reported. Thus, for the present comparisons we aimed to observe the qualitative metabolite changes within the experimental and simulated data, noting complications in making rigorous quantitative comparisons.

F16BP and PEP were measured reported as individual metabolites in the two cancer cell lines, A2780 and HCT116. The comparison of the simulated and experimental data is presented in Figs 6-7 and 10-11. Because of the difficulties in distinguishing isobaric metabolites from one another, G6P+F6P and GAP+DHAP were grouped in the experimental data, and the sum of these metabolites are reported. The model is able to determine the individual metabolite concentrations, however, following each simulation the two metabolites from the model data (G6P+F6P and GAP+DHAP) were added for a closer comparison to the experimental data. Moreover, the data was normalized so that each experiment (*in vitro* and *in silico*) began with the same metabolite concentration, again for a clearer comparison of the actual changes occuring as FK866 doses increased (experimental) and as GAPDH inhibition increased (model simulations).

*In silico*, we observed increases in all upper glycolytic metabolites with inhibition of GAPDH, supporting the metabolic data of NAMPT inhibition. The lower glycolytic metabolite, PEP shows reduction following increased inhibition of GAPDH, in agreement the results seen in the experimental data. Using a K/P ratio of 0.1, F26BP was the only metabolite that did not change mean values over time throughout the course of GAPDH inhibition at any level. Notably, the experimental data shows a fairly wide range of metabolite concentrations with similar inhibitor doses, between and even within both cell lines. For example, the G6P+F6P concentration in the HCT116 cell line increased from 0.052 mM to 0.512 mM with the highest FK866 treatment, and in the two separate experiments with the A2780 cells the metabolite concentrations increased to only 0.18 mM and 0.13 mM (Fig 4). Again, the aim for the inhibition simulations was to observe the overall trend of metabolite changes, meant for a qualitative comparison. There are slight variations from the experimental data and the model data. Still the results are similar or within range of the experimental data; in the model simulation the F16BP concentration increased from 0.0022 mM to 0.0548 mM at the highest inhibition level, while the F16BP concentration in the HCT116 cell line rose from 0.0022 mM to 0.0496 mM at the highest FK866 treatment (Figs 6-7). Specific kinetics and prior knowledge of experimental data may only aid in reproducing even more consistent results in the future.

The glycolysis model was additionally simulated in the SimBiology app in MatLab using the ode15s solver [32]. The rate equations, parameters and initial concentrations were identical to those used in the queueing theory model in the steady state simulation. When running the simulations, the metabolites reached steady state levels up to 10 seconds. Following the 10 second mark, however 2PG and 3PG progressed to positive infinity and negative infinity, respectively. Moreover, many metabolites had negative concentration values in the steady state simulations. The compiled SimBiology ode15s solver was easy to both operate and simulate deterministic outcomes quickly, however the simulation produced unstable and negative values. As mentioned in the methods section, queueing theory modeling ensures positive concentrations, a clear benefit when attempting to track metabolite or biological species concentration changes. In the uncompiled version, the current queueing theory model was able to complete 20 minutes of simulation time for 30 cells in roughly two hours.

## Conclusion

The paper presents a pathway model of glycolysis as a queueing network, a modeling approach widely used in modeling telecommunication packet networks. Dynamic modeling of biological systems, while exceptionally useful poses certain limitations computationally and in reproducing observed phenotypic changes. The application of queueing theory in dynamical modeling may offer a method to overcome such limitations. The current applications of this work hold significant promise for advancing computational biology and biochemical research. The queueing theory represents a mainstay modeling approach of telecommunication networks with application to simulate intracellular metabolism. By viewing enzymes as “gates” and their substrates as “packets,” we have reduced the computational complexity of the simulation to the advantage of much more rapid calculation. Moreover, we have shown previously that we can model intracellular mechanisms and do so while simulating the random variation which exists between and within living cells.

Research and experimental techniques in metabolomics have rapidly evolved since its introduction. Modeling strategies must also be able to be adaptable to accommodate novel information and amend the data as needed. The modularity of queues incorporated provides a suitable approach for further model extension, whether that be additional metabolic reactions, parameter refinement, or multi-scale modelling approaches. Moreover, this approach enables the ability to simulate biochemical reactions stochastically without the need to implement or solve stochastic algorithms. As seen above, GAP and DHAP were represented experimentally as a combination of metabolic intermediates, due to their chemical similarities. Although MS technological methods have become increasingly sensitive to detecting small molecules, isobaric metabolites are often difficult to distinguish from one another. This is the case not only for several metabolic intermediates of glycolysis, but also to additional metabolic pathways. An advantage to the in silico mechanistic modeling of metabolic networks, is the ability to represent such metabolites as individual entities investigating distinct metabolic reactions and the dynamics of each metabolite providing a more in-depth observation of the intracellular interactions.

The need for models to be informed from and then simulate data using metabolomics sources represents a significant advance in future possibilities of the use of this approach. With the small-scale investigation and advanced and large-scale experimental biology, computers have become pivotal in not only managing data but also in understanding the biological significance of the results and developing further hypotheses for future research. Due to the ability to change variables and quickly analyze the resulting metabolic effects, investigators can then simulate the effects of drugs or mutation on such processes. In all, the ability to accurately and quickly simulate intracellular and intra-tissue pathways represents a considerable leap forward in the ability to understand central biochemical underpinnings of cellular life. The advancement of technology in both experimental biology and computational systems has allowed scientific discovery and investigation on the chemical level. Elucidation of intracellular metabolite and chemical dynamics can provide valuable insight into how cells utilize cellular components to grow, respond to environmental stimuli, and ultimately support life. While deterministic models are able to describe glucose metabolism and metabolic systems in general, we believe queueing theory may have the potential to more realistically describe metabolic behavior by providing stochasticity to the pathway. In summary, the current study presents the application of queuing theory as a beneficial modeling approach in simulating metabolic pathway dynamics and predicting the effects of pharmacological inhibition.

## Acknowledgments

The following support is acknowledged: NIH GM103427 (PHD), and the University of Nebraska at Omaha Office of Research and Creative Activities FUSE and GRACA program (EC). This work utilized the Holland Computing Center of the University of Nebraska, which receives support from the Nebraska Research Initiative.

## Supporting Information

**S1. Pseudo Code.**

**S2. Kinetic Constants and Parameters of the model.**

**S3. Rate Equations used in the model.**

**S4. Maximal Velocities.**

**S5. Rate Files**

